# Tether-mediated extraction of myelinoid bodies by microglia and astrocytes can maintain myelin integrity

**DOI:** 10.64898/2025.12.05.692364

**Authors:** Erik Späte, Kieran P. Higgins, Yeonsu Kim, Carolina Mangana, Lina Komarek, Mohammadhossein Teymouri, Ulrike C. Gerwig, Svea Melzer, Boguslawa Sadowski, Torben Ruhwedel, Predrag A. Janjic, Bastiaan Lodder, Andrew J. Dwork, Maarten H.P. Kole, Anna M. Steyer, Wiebke Möbius, Klaus-Armin Nave, Sandra Goebbels

## Abstract

Oligodendrocytes make myelin for the electrical insulation of axons and saltatory impulse conduction. Myelin lipids and proteins undergo a slow turnover, but exactly how the multilamellar and compacted membrane sheaths are remodeled without compromising myelin sheath integrity has remained puzzling, in particular at advanced age when myelin abnormalities increase. Earlier EM studies had suggested myelin membranes are shed and subsequently phagocytosed by microglia. However, the formation of multilamellar myelinoid bodies (MBs), leaving a well-ordered myelin sheath behind, is difficult to reconcile with simple shedding mechanisms. Here, we show by three-dimensional FIB-SEM reconstructions of optic nerves in mice and by two-photon live-imaging of myelinated cortical slices that MBs are initially connected to their parental sheaths by long tethers, which are stretched by trogocytosing microglia and astrocytes. We observe ruptured tethers attached to both MBs and sheaths, suggesting a novel mechanism of tension-driven tether scission. Importantly, the successive fusion of the corresponding innermost myelin membranes in an extended tether can preserve myelin sheath integrity. Thus, the remodeling by tether-mediated MB extraction emerges as a mechanism of physiological maintenance of myelin sheaths in the CNS.

## Introduction

In the vertebrate central nervous system (CNS), oligodendrocytes grow lipid-rich myelin membranes around associated axon segments, elaborating tightly wrapped, remarkably stable myelin sheaths for electrical insulation and efficient neuronal signal propagation (*1*). In the brain, most oligodendrocytes are formed during childhood and persist throughout adult life (*2*). While myelin structural proteins and lipids are among the longest-lived in the body, they are replaced more rapidly than the sheath itself (*3, 4*). This requires continuous renewal of myelin membranes, involving the turnover and incorporation of myelin proteins and lipids, first suggested by metabolic labelling studies (*5*).

Understanding the mechanisms underlying myelin turnover in the aging brain has become increasingly important as age-dependent myelin structural defects have emerged as drivers of neurodegenerative disorders, such as Alzheimer’s and Parkinson’s disease (*6–8*). Most progress has been made in understanding macrophage/microglia-mediated myelin breakdown and myelin debris removal in demyelinating diseases. Here, myelin splitting, cutting, and finally peeling of myelin lamellae by activated macrophages/microglia, resulting in complete destruction and phagocytosis of the myelin sheath, is well documented (*9–13*). In demyelinating diseases, cortical ischemia, and spinal cord injury, myelin debris has also been detected inside astrocytes, which are capable, like microglia, of phagocytosing myelin *in vitro* (*14–16*). Finally, the engulfment and removal of entire myelin sheaths by microglia has been observed during development in the zebrafish (*17*) suggesting a role for microglia in the pruning of oligodendrocytes and their processes that are developmentally produced in excess.

On the other hand, little is known about the mechanisms underlying physiological myelin turnover, including the relative contributions of astrocytes, particularly in the aging brain, where the need to clear defective myelin increases. Although myelin aging is marked by significant structural alterations, such as ballooning, outfoldings of redundant myelin, and splits of the major dense line (*18*), the disease-associated pattern of macrophage/microglia -mediated myelin breakdown, as described for demyelinating mouse models (*10*), has not been reported in aging. Nevertheless, multilamellar globular myelin breakdown products, termed myelinoid bodies (MBs), accumulate both inside cells and in the extracellular space (*5, 19–21*). This raises the possibility that aging involves a distinct myelin turnover/clearance mode. Indeed, based on immunofluorescence microscopy and two-dimensional EM analyses, it has been suggested that MBs are shed from myelin sheaths into the extracellular space, where they are subsequently engulfed and degraded by microglia (*21, 22*). However, exactly how MBs are detached from intact sheaths without damaging the remaining myelin membranes remained unresolved.

In the present study, we employed 3D volume electron microscopy and 2-photon live-cell imaging and discovered that only a small fraction of extracellularly located MBs are completely detached from the myelin sheath, indicating that shedding is not the primary mechanism of myelin removal. Instead, by a mechanism reminiscent of trogocytosis, microglia and astrocytes actively extract several outer myelin layers forming a multilayered MB that innitially remains connected to the sheath by a thin tubular myelin tether. We propose a novel model, in which the stretching of this myelin tether over several micrometers allows for the consecutive inner-lamellae fusions within the tether, which gradually re-seals the myelin lamellae and allows for the complete uptake of the MB after tether rupture. Our results indicate that under physiological conditions in the adult, and increasingly during aging, the uptake of myelin debris is an active phagocyte-driven process, rather than a passive consequence of myelin fracturing or shedding. We also reveal a prominent role of astrocytes in aging-associated myelin turnover when microglia are increasingly burdened by myelin degradation (*23*).

## Results

### Aberrant Myelin Ultrastructure in the Aging Optic Nerve

To investigate pathological abnormalities that may contribute to the increased degradation and turnover of myelin in the aging CNS, we examined chemically fixed and high-pressure frozen (HPF) mouse optic nerves, comparing adult mice at age 6 months (mo) with mice of advanced age (24 mo). Cross-sectional analyses were conducted using transmission electron microscopy (TEM, **Fig. 1a**). Aging was associated with significant increases in both axonal calibers and the average number of myelin sheath lamellae (**Fig. 1b**). Small calibers axons with diameters of 0.8 µm or less exhibited a genuine increase in myelin sheath thickness with age (**Fig. 1b**). Myelinated axons exceeding 0.8 µm in diameter maintained normal myelination, with additional myelin layers matching increased axonal calibers with age (**Fig. 1b**).

**Figure 1:**
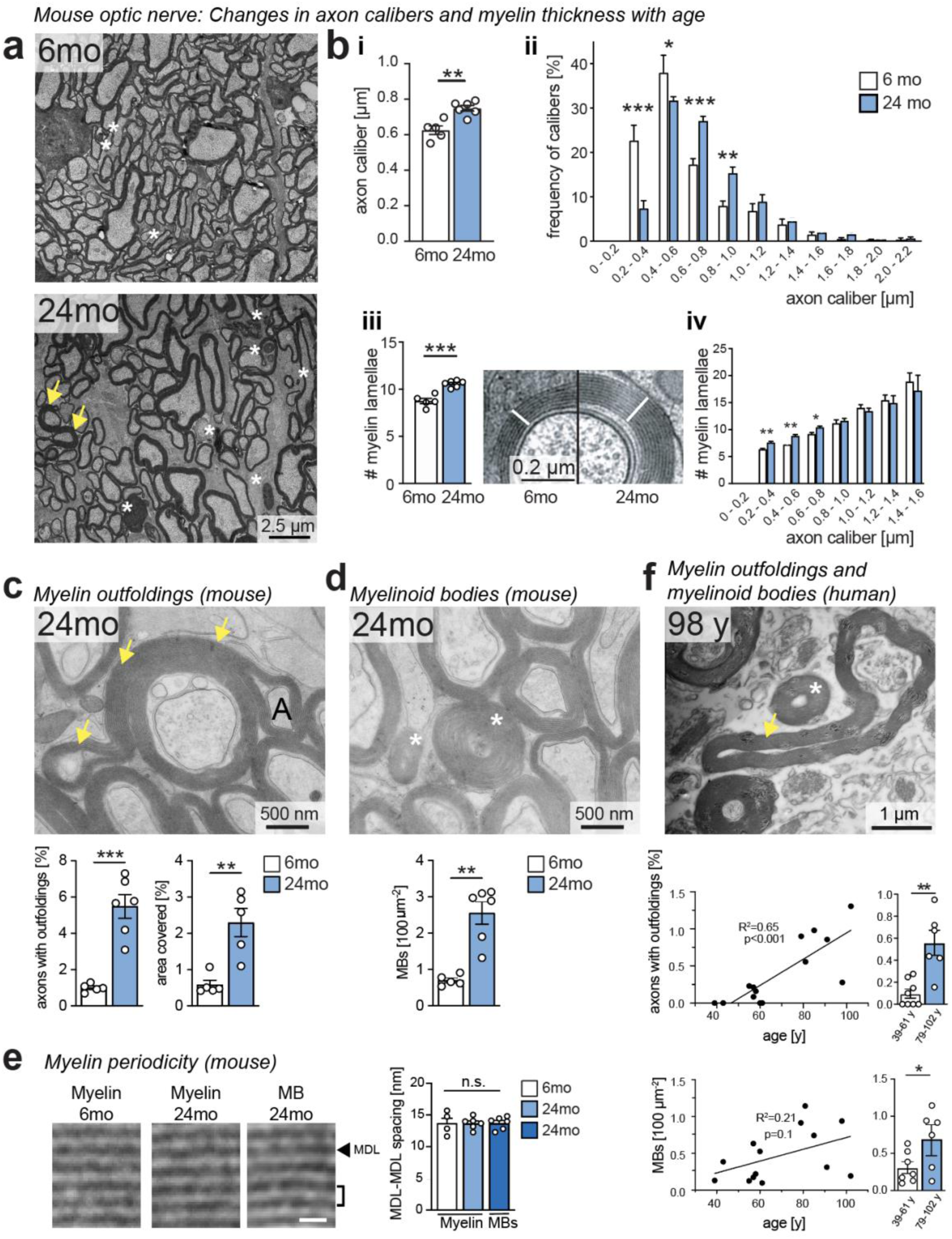
Myelin outfoldings and MBs increase with aging in mouse and human optic nerves. a) Representative cross sections of chemically fixed and epon-embedded optic nerves of wildtype mice at ages 6 mo (top) and 24 mo (bottom). Asterisks indicate myelin debris in form of MBs, arrows point to myelin outfolding. b) Quantitative assessment on HPF optic nerve samples shows a significant shift of axon calibres towards larger diameters with age (i, ii) (left: diff_6mo-24mo_ = 0.12 ± 0.03 µm, Student’s t-test with Welch’s correction; right: F(9, 90) = 12.31, 7.91% of variation, two-way ANOVA with Šidák correction). Accordingly, the average number of myelin lamellae is increased with age (diff_6mo-24mo_ = 1.83 ± 0.30, Student’s t-test with Welch’s correction) (iii), resulting in normal myelin sheath thickness for axons above 0.8 µm in diameter and a slight hypermyelination of axons below 0.8 µm in diameter in aged optic nerves (iv) (diff_0.2-0.4_=1.59 ± 0.34, diff_0.4-0.6_=1.24 ± 0.28, diff_0.6-0.8_=1.24 ± 0.37, multiple t-test with Holm-Šidák correction). c) Quantitative analysis of myelin outfoldings (arrows in upper panel) on electron microscopic images of HPF mouse optic nerves samples at ages 6 mo and 24 mo. Number and area of outfoldings significantly increases with aging (left: diff_6mo-24mo_ = 4.49 ± 0.72 %; right: diff_6mo-24mo_ = 1.71 ± 0.41 %, Student’s t-test with Welch’s correction). d) Number of myelinoid bodies (MB, asterisks in upper panel) that appear unconnected to myelinated axons significantly increases with aging (diff_6mo-24mo_ = 1.85 ± 0.35 per 100 µm^2^, Student’s t-test). e) Representative images and quantitative analysis of myelin periodicity in 6 mo and 24 mo myelin and in 24 mo well-preserved MBs in HPF optic nerve samples shows no significant difference (n=4-6, myelin_6mo_ = 13.7 ± 1.82 nm, myelin_24mo_ = 13.7 ± 0.93 nm, MB_24mo_ = 13.7 ± 0.98 nm, F(2, 13) = 0.002, one-way ANOVA with Sidak’s multiple comparison test). Scale bar = 15 nm; arrow head, major dense line (MDL). f) Quantitative analysis on electron microscopic images of chemically fixed epon-embedded human optic nerves (representative image on top). Percentage of axons with outfoldings (arrow in upper panel) and MB (asterisks in upper panel) density is significantly increased in human optic nerves during aging (outfoldings left: slope = 0.017 ± 0.004 %/year, linear regression; right: diff_39-61y–79-102y_ = 0.73 ± 0.13 %, Student’s t-test with Welch’s correction; MBs left: slope = 0.001 ± 0.005 %/year, linear regression; right: diff_39-61y–79-102y_ = 0.39 ± 0.17 %, Student’s t-test with Welch’s correction). n_mouse_ = 5-6, n_human_ = 6-8; 1000 axons analyzed per individual; t-tests are unpaired and two-tailed, slope significance was estimated via F-Test; errors are SEM; * p < 0.05, ** p < 0.01, *** p < 0.001.

Several known age-related myelin abnormalities, such as thin myelin and decompacted and split sheaths (*18, 19*) were noted in aged optic nerves and comma-shaped outfoldings were prominent (**Fig. 1c**). Viewed in cross section, these outfoldings often coiled around adjacent myelinated axons and increased with age both in number and size (**Fig. 1c**). Additionally, myelinoid bodies (MBs) were increased in numbers with age (**Fig. 1d**), often composed of multilamellar membrane stacks with the normal periodicity of myelin when analyzed by EM in high pressure-frozen (HPF) material (**Fig. 1e**). On EM cross sections, MBs appeared frequently close but not connected to nearby myelin sheaths (**Fig. 1d**). By comparing cross-sections of Epon-embedded optic nerves of human autopsies at middle age (39-61 years (y)) and old age (79-102 y), both MBs and comma-shaped outfoldings also increased in number, the latter generally shorter in humans than in mice (**Fig. 1f**). We conclude that MBs and outfoldings are a feature of aging in both human and mouse white matter.

Next, we sought to localize the most recently formed MBs in the mouse, i.e. prior to enzymatic degradation, using antibodies against the myelin protein PLP and the astroglial marker protein GFAP with subsequent immunogold EM labeling. In 24 mo old optic nerves, PLP positive MBs (immunolabeled by 10 nm gold particles) localized in non-compact myelin compartments, (**Fig. 2a, Suppl. Fig. 1a**), in GFAP-negative microglia (**Fig. 2b**), in astrocytes (**Fig. 2c**) and in the extracellular space (**Fig. 2b-c)**. Microglia, which were identified by their cytoplasmic dark bodies (lysosomes), and a small nucleus with dense chromatin, were characterized by inclusions that are presumably remnants of lysosomal myelin degradation. These appeared as multilamellated lipid-rich membrane stacks with closely associated cholesterol crystals and occasional membrane fusion profiles (**Fig. 2b**), but largely without remaining PLP immunoreactivity. The identification of PLP-positive MBs along with occasional findings of PLP-positive lysosomes in aged GFAP immuno-labeled astrocytes (15 nm gold particles) (**Fig. 2c**), indicated that astrocytes contribute to the clearance of myelin debris in the aging optic nerve.

**Figure 2:**
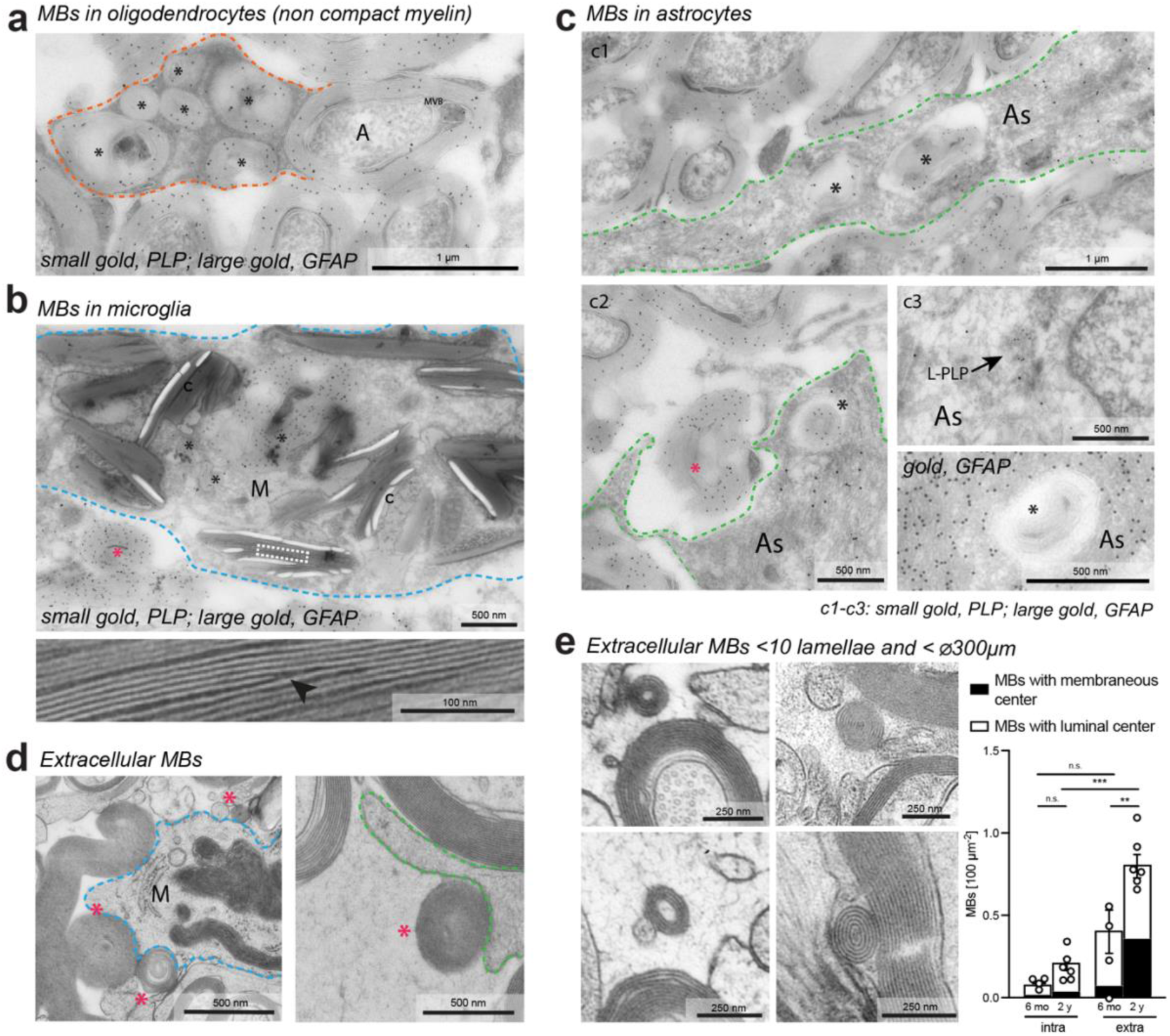
Localization of MBs. a)-c) Colloidal immunogold-EM labelling for GFAP (15 nm gold-particles) on ultrathin optic nerve sections of wildtype mice at age 24 mo. MBs (co-labelled for PLP by 10 nm gold-particles in a, b, and c1-c3) localize extracellularly (mangenta asterisks) and intracellularly (black asterisks) a) in a non-compact outer tongue (outlined in orange) of the myelin sheath around an axon (A), b) in a microglial cell (M, outlined in blue), and c) in an astrocyte (As, outlined in green). Microglia are characterized by parallel membrane stacks depicting occasional fusion/fission profiles (lower panel in b, arrowhead) that are associated with cholesterol crystals (c) and likely represent myelin debris at advanced stage of degradation. GFAP-positive astrocytes occasionally contain PLP-positive lysosomes (arrow in c3, L-PLP), indicating their active involvement in the degradation of myelin debris. d) HPF EM images depict differently sized, extracellular MBs (mangenta asterisks) in close association to microglia (left panel, outlined in blue) and astrocytes (right panel, outlined in green). e) Tiny, well-layered MBs (<10 lamellae and < 300 nm in diameter) with either luminal (electron micrographs on the left) or membranous center (electron micrographs on the right) are predominantly localized extracellularly and increase in abundance with age. Two-way ANOVA followed by Šidák multiple comparison test on total density of MBs. Data are represented as mean ± SEM. Individual data points represent total density of MBs of each animal (n = 4-6). MVB, Multivesicular Body.

PLP-positive MBs in the extracellular space often located near phagocytes (**Fig. 2b-d**). Detailed analyses of well-preserved HPF samples revealed many MBs to be < 0.3 µm in diameter with often few myelin lamellae (**Fig. 2e**). These ‘tiny’ MBs still had a regular periodicity and either an electron-light core (termed ‘luminal’ center) or an electron-dense core with membrane-profiles (termed ‘membraneous’ center) (**Fig. 2e**). ‘Tiny’ MBs were more often located extracellular than intracellular (p < 0.001), had more often a luminal than membranous center (p = 0.003), and were increased in numbers at age 2 years (p = 0.001, three-way ANOVA). The largest age-dependent increase was seen in ‘tiny’ extracellular MBs with a membranous core (p = 0.022, Šídák’s multiple comparisons test) (**Fig. 2e**). This raised the question about the origin of these peculiar extracellular structures.

### Myelinoid bodies outside of oligodendrocytes are commonly tethered in the aged optic nerve

To better understand age related myelin abnormalities, we employed focused ion-beam scanning electron microscopy (FIB-SEM) of myelinated optic nerves at 6 and 20 mo of age and acquired volumetric stacks for subsequent 3D reconstructions.

As expected myelin outfoldings were particularly evident in the aged optic nerve. While appearing comma-shaped in 2D transverse sections, they emerged in 3D as flat sheets of compact myelin extending predominantly longitudinally along the myelinated axon with an average sheet-length of 9 µm (**Fig. 3a, Suppl. Movie 1**). A cytoplasmatic compartment, continuous with the inner tongue, lined the edges of the outfoldings. Their sheet-like structure restricts membrane bending along the Z-dimension, which explains why in 2D EM images myelin outfoldings generally project perpendicularly from the cross-sectioned axon.

**Figure 3:**
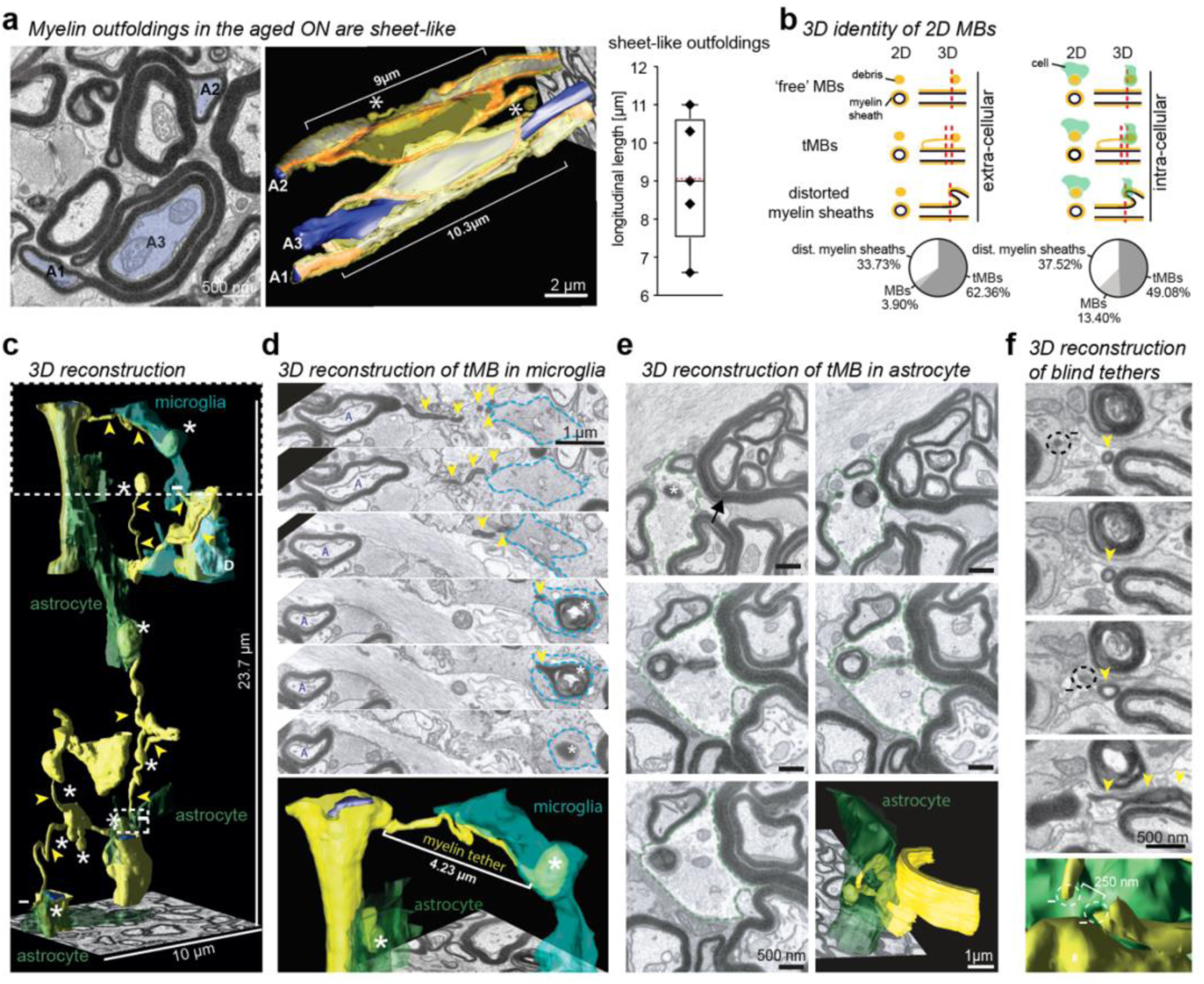
MBs engulfed by phagocytes are connected to myelin sheaths and myelin outfoldings by thin tethers. 3943 focused ion beam-scanning electron microscopy (FIB-SEM) micrographs were scanned from a tissue block of optic nerve (age 20 mo) to cover a volume of ≥15 μm × ≥ 15 μm × ≥ 197 μm with a voxel size 2 nm x 2 nm x 50 nm and were subsquently used (binned by 5 in x/y) for the analyses of myelin outfoldings and MBs. a) SEM micrograph and three-dimensional (3D) reconstruction of myelin sheaths and myelin outfoldings (colored in yellow for compact myelin and orange for non-compact myelin) and the respective three myelinated axon segments (A1-A3, colored in blue). Myelin outfoldings that appear “finger-like” in 2D (left panel) mostly represent sheets in 3D view with an average length of 9 µm (based on n=5 reconstructed outfoldings). Along the entire edge of the outfolding stretches a cytoplasmatic compartment that is directly connected to the inner tongue. Two MBs (asterisks) are connected to the myelin sheath and the outfolding of A2, respectively by a short myelin tether. b) Classification and quantitation of extracellular (left) and intracellular (right) located MBs in phagocytes based on 1200 consecutive FIB-SEM images. MBs that are identified as allegedly unconnected in 2D can be in one of three categories in 3D: truly unconnected ‘free’ MBs, MBs that are still tethered to myelin (tMB), and miscategorized MBs that are the consequence of distorted myelin sheaths or myelinated axons. tMBs are the largest category both intracellularly and extracellularly. Red dashed lines indicate section planes that all present similar in 2D view. c) 3D reconstruction of myelinated axon segments (axon, blue; myelin, yellow), myelin debris including tethers and tMBs (yellow), associated astrocytes (identified by their characteristic parallel intermediate filaments, green), and microglia (identified by multilamellar debris and cholesterol crystals, blue-green). Arrowheads, myelin tethers with associated MBs; -, blind-ending myelin tethers without associated MB; asterisks, MB; D, myelin debris in microglia at advanced degradative state. d) Magnification from c). A tMB (asterisk) surrounded by a microglia process that is still connected to a myelin sheath (yellow) by a thin myelin tether (yellow, arrowheads). A, axon. e) 3D reconstruction of an astroglialy located tMB (asterisk) that is still tethered to the edge of a myelin outfolding (arrow). f) Magnification from c), lower part. Two ‘blind’ tether ends in close proximity, with one connected to a myelin sheath and the other one connected to an MB located in an astrocyte.

Moreover, 3D reconstructions in the aged optic nerve revealed that many MBs, which usually appear detached in 2D EM, are in fact still continuous with adjacent myelin sheaths. Occasionally, MBs located within non-compact myelin, such as swollen inner tongues, were still connected to the sheath via a thin and short myelin neck (**Suppl. Fig. 1**), consistent with a model, in which those MBs might facilitate cell-autonomous myelin degradation in oligodendrocytes (*24, 25*). Strikingly, even the more common MBs located extracellularly or engulfed by phagocytes, which are thought to clear most myelin debris via lysosomal degradation (*21, 26*), frequently retained a thin myelin tether traceable to a parental sheath, also when positioned several micrometers from any myelinated axon (**Fig. 3**).

Thus, we next quantified different categories of MBs located extracellularly or within phagocytes by analyzing 1200 serial 2D images from the 20 mo FIB-SEM stack. In each 2D image, we identified all MBs^2D^ and classified them based on their 3D identity as: (a) fully separated from a myelin sheath (= ‘free‘ MBs), (b) still connected to a parental sheath by a thin myelin membrane tether, or (c) unrelated myelin distortions that appeared circular in 2D (**Fig. 3b**). Surprisingly, only 3.9% of the extracellular MBs^2D^ were truly separated from their parental myelin sheath in 3D view. In the same volume, 13.4% of the ‘intracellular’ MBs^2D^ that localized to astrocytes or microglia were fully separated (**Fig. 3b, Supp. Fig. 2a**). An additional 33.7% of extracellular MBs^2D^ (and 37.5% of intracellular MBs^2D^) were myelin distortions, caused by local bulging, protrusions, delaminations, or axonal sprouts (**Fig. 3b,c Supp. Fig. 2b**). However, the vast majority of MBs^2D^ (intracellular MBs^2D^: 62.4%; extracellular MBs^2D^: 49.1%) were connected by a thin myelin tether to a parental myelin sheath. We therefore termed these structures ‘tethered’ MBs (tMBs) (**Fig. 3b,c; Supp. Fig. 2c**) and conclude that the majority of MBs in 2D images are in fact cross-sectioned myelin tethers or MBs at the distal ends of these tethers rather than shed myelin debris. We note that myelin tethers are, even when oriented perpendicular to the axon orientation, clearly distinguishable from myelin outfoldings, because they are comprised only of superficial myelin lamellae, whereas outfoldings are caused by the outbulging of entire myelin sheaths in their full thickness.

3D reconstructions indicated that tMBs stretch away from intact myelin sheaths, or the lateral edges or tips of myelin outfoldings as well as from myelin distortions (**Fig. 3a, c, d, e**, **Fig. 4, Suppl. Movie 1, 2**). Myelin tethers varied in length from short structures of <1 µm to longer tethers exceeding 24 µm. Their spherical MBs were mostly but not always engulfed by a microglial or an astroglial process (**Fig. 3d-e, Suppl. Movies 3, 4**). Notably, while enriched with aging, tMBs were also occasionally detectable in the 6 mo FIB-SEM dataset (**Suppl. Movie 5**), and thus not exclusive to aging.

**Figure 4:**
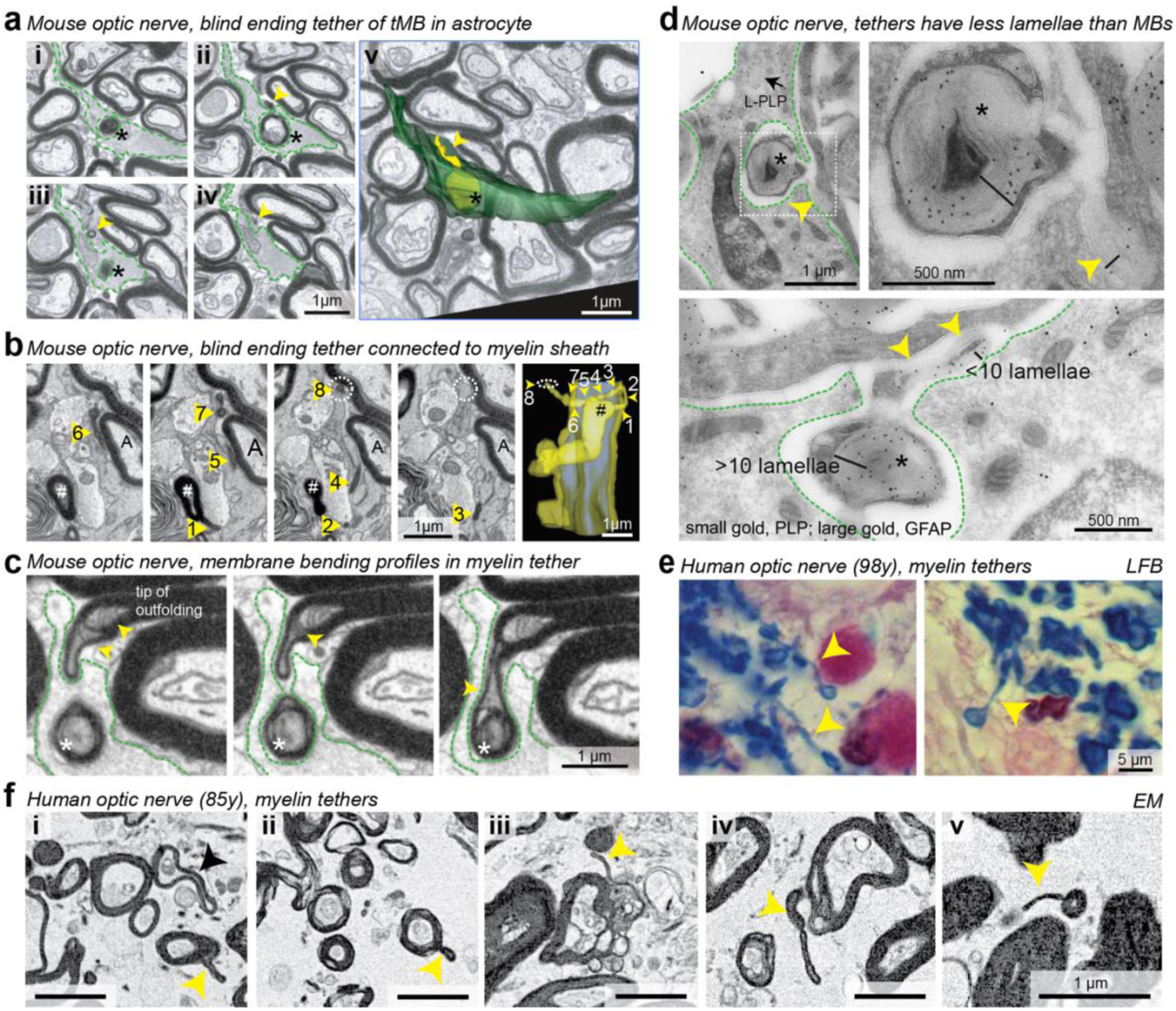
Myelin tethers thin out and rupture. a-c: FIB-SEM analysis and 3D reconstructions of mouse optic nerve (age, 20 mo). a) An astrocyte (green) engulfs an MB (asterisk), of which a short tether (arrowhead) sticks out that is not connected to a myelin sheath. b) A thin blind ending myelin tether (labelled by arrowheads 1-8 that correspond to respective numbers in the electron micrographs selected from the FIB-SEM image stack) is connected to the myelin sheath (yellow) of an axon (A) via a thicker finger-like myelin protrusion (yellow, #). c) A tMB (asterisk) engulfed by an astrocyte (outlined by green dashed line) with its tether connected to the tip of an outfolding. Bendig membrane profiles (arrowheads) are compatible with intermembrane fusion and fission of inner tether membranes. d) 2D immunogold labelled images show two astrocytes (outlined by green dashed lines; top left and bottom panel) engulfing tMBs (top right panel, higher magnification). Tethers (arrowheads) are comprised of less myelin lamellae than the corresponding MB. Arrow indicates a lysosome containing PLP-positive material (L-PLP) e) Luxol Fast Blue (LFB) staining of an aged human optic nerve shows myelin tethers (yellow arrowheads) reminiscent of myelin tethers found in aged mouse optic nerves. f) SEM analysis of aged human optic nerve shows examples of thin myelin tethers associated with myelin sheaths (i-ii), with myelin dystrophies (iii-iv), or connected to an MB (v). Note that myelin tethers (yellow arrowheads) are distinguishable from myelin outfoldings (black arrowhead).

Moreover, 3D reconstructions in the 20 mo-old optic nerve revealed that myelinated axon segments occasionally appeared bent outward toward the associated tether, indicative of a pulling force exerted by the phagocyte that had engulfed the associated tMB (**Fig. 3d**). Additionally, we observed blind-ending tethers (**Fig. 3f**) and identified both tMBs with tethered endings protruding from phagocyte’s cups (**Fig. 4a**), as well as tethers originating from a myelin sheath but not connecting to any MB (**Fig. 4b**), indicating prior tether rupture. 3D reconstructions also showed that inner myelin lamellae within intact tethers curved back on themselves to form loop-like bending profiles, while the outer lamellae remained continuous (**Fig. 4c**). Such configurations are consistent with membrane apposition and fusion among the innermost layers, resulting in progressive tether thinning. In agreement, tethers of tMBs contained fewer myelin layers than the corresponding MBs **(Fig. 4d**). Consequently, we suggest that “tiny” MBs with few layers and central membrane profiles observed in 2D (**Fig. 2e**) in fact represent cross-sectioned tethers with inner lamella fusion.

Light microscopic analyses of LFB-stained human optic nerves and subsequent electron microscopic analyses indicated that tMBs and myelin sheaths, associated by myelin tethers, are also present in aged human optic nerves (**Fig. 4e, f**).

Taken together, our ultrastructural results indicate that MBs are not simply released from myelinated axons but that both astrocytes and microglia actively stretch tMBs as a living part of the outer myelin sheath. We propose that sequential fusion of the innermost tether membranes results first in thinning and finally in rupture of the tether, after which the remaining ends retract, allowing full internalization of the tMB. This mechanism would represent an adaptation of trogocytosis, the ‘nibbling’ of small parts of the membrane from another cell (*27*).

### Microglia and Astrocytes Clear Myelin Debris in the Brain

Next, we confirmed that both microglia and astrocytes in the aged brain exhibit increased uptake of myelin debris. We performed co-immunostaining on paraffin sections from mouse brains aged 6 mo and 24 mo, using antibodies against myelin-basic protein (MBP), IBA1, and GFAP to label myelin, microglia, and astrocytes, respectively (**Fig. 5a,b**). At age 6 mo, 21,6% ± 2,4% of microglia and 9,7% ± 5,5% of astrocytes contained MBP-positive myelin debris when quantified in the stratum lacunosum moleculare (SLM) of the hippocampus. At age 24 mo, these numbers increased to 56% ± 2,5% for microglia and 39,3%± 3,8% for astrocytes (**Fig. 5c**). At both ages myelin debris inclusions were approximately ten times larger in microglia than in astrocytes (**Fig. 5c**). Although we confirmed the increased uptake of myelin debris by astrocytes at advanced age, the fraction of astrocytes containing myelin debris at age 6 mo was low (**Fig. 5c**). Hence, when we next investigated the mechanisms of myelin tether formation in an *in vitro* model of cortical organotypic slice cultures that corresponds to young mouse age, we focused on microglia.

**Figure 5:**
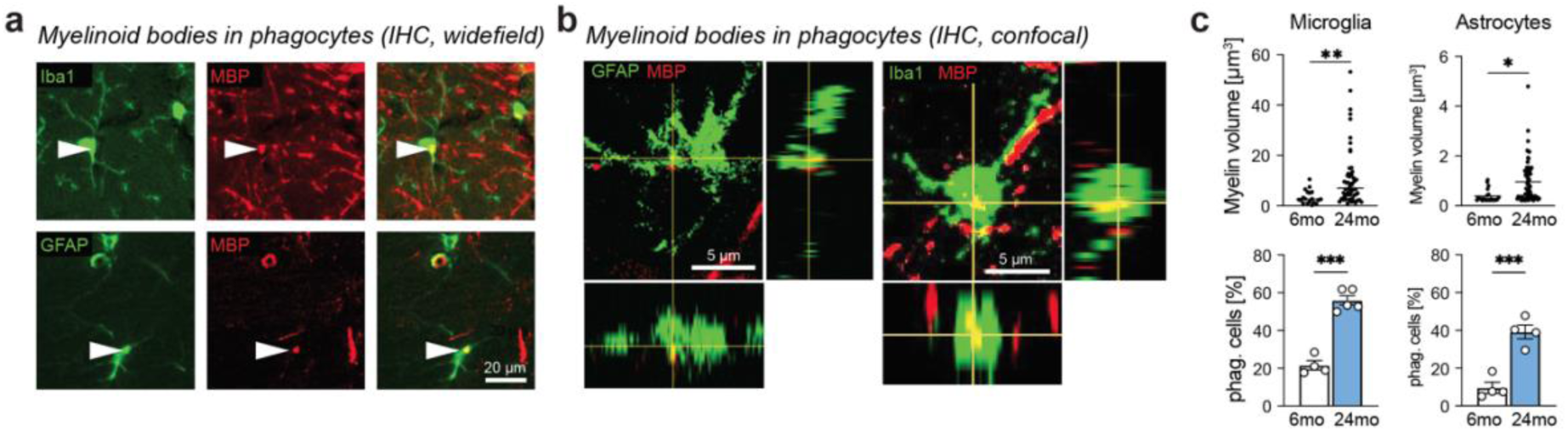
Microglia and astrocytes clear away myelin debris in the brain. a) Representative images for presence MBP positive (red) myelin debris (arrowhead) in anti-GFAP immuno-labelled astrocytes (green) and in anti-Iba1 immunolabelled microglia (green) in aged (24 mo) mouse stratum lacunosum moleculare (SLM) of the hippocampus. b) Confocal 3D projections of a representative astrocyte (left, green) and a microglia (right, green) showing MBP positive inclusions (red). c) Quantification of the volume of myelin debris within microglia and astrocytes (top) and quantification of the percentage of astrocytes and microglia that contain myelin debris (bottom) within the SLM of the hippocampus at ages 6 and 24 mo. Upper: Data points represent individual myelin inclusions with median, n = 3-6 mice, Linear Mixed Model (see methods); Lower: Data represented as mean +/- SEM, n=4-5 mice. Student’s t-test; *p<0.05. **p<0.01, ***p<0.001.

### Microglia Actively Stretch tMBs in Cortical Organotypic Slice Cultures

Organotypic slice cultures undergo similar key developmental steps as observed in brain development *in vivo*, including myelin formation, and have been used to study microglial functions (*28, 29*). To study the dynamics of microglial interactions with myelin sheaths, we prepared cortical organotypic slice cultures and virally transduced them with AAV-Mbp:memEGFP to express a membrane-linked EGFP in oligodendrocytes. We first fixed and stained the slice cultures for MBP, Iba1, and EGFP to verify myelin coverage across the slice at DIV 21 (**Fig 6a**). Confocal imaging confirmed sparse and selective labelling of oligodendrocytes as previously described (**Fig 6b**) (*30*). In addition, TEM analysis revealed myelinated axons with normal g-ratio and typical myelin ultrastructure including compact myelin and non-compact inner and outer tongues (**Fig 6c**).

**Figure 6.**
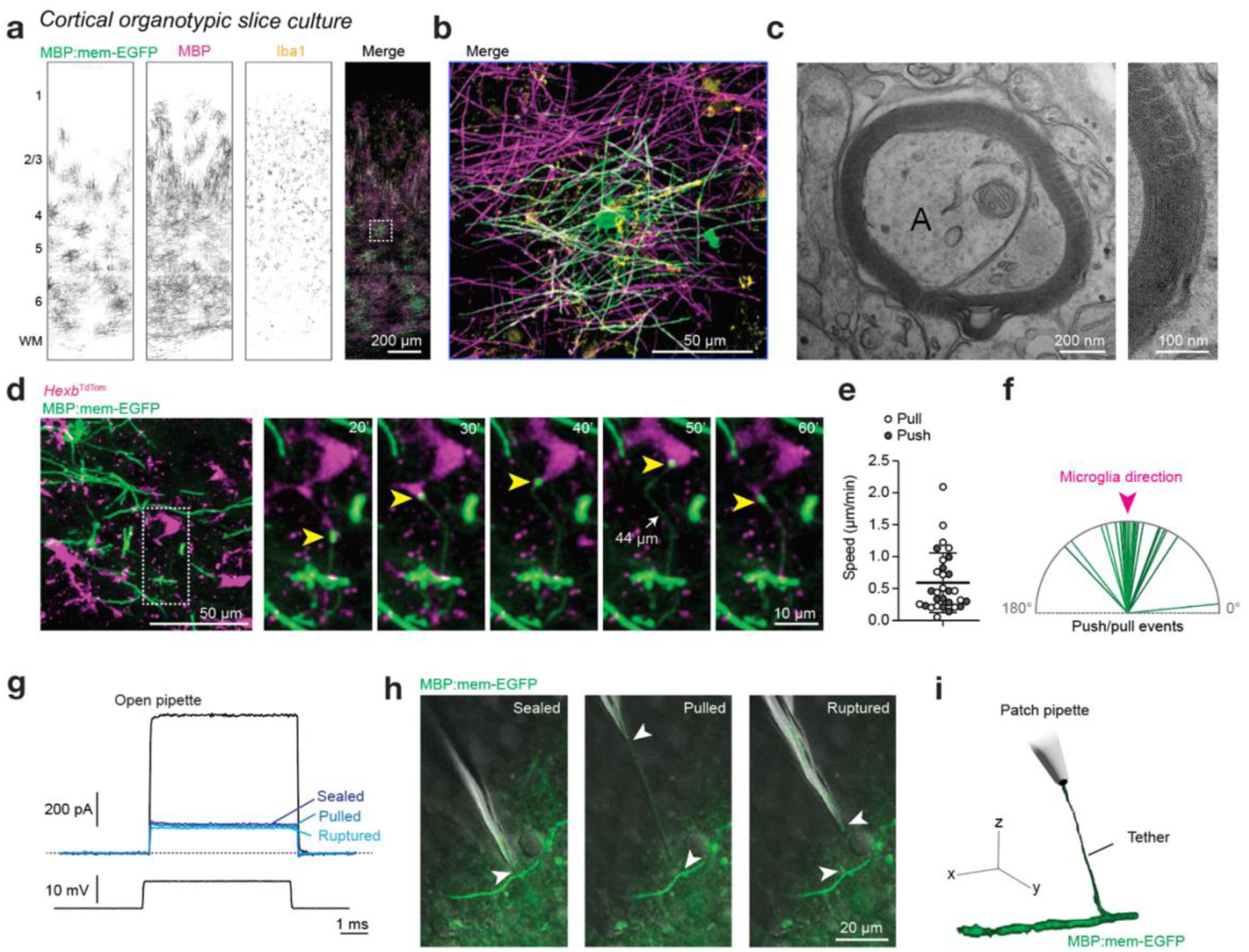
Microglia actively stretch tMBs in cortical organotypic slice cultures. a) Cortical organotypic slice cultures showing robust myelination at DIV 21. Representative confocal images of a slice stained for AAV-MBP:mem-EGFP (green), MBP (magenta), and Iba1 (yellow). b) Magnification of a single oligodendrocyte shown in a (white) box. c) Left, Representative TEM image of a myelinated axon (A, diameter 704 nm) with stereotypic ultrastructure of compact and noncompact myelin. Right, zoom in showing the compact layering of 10 lamellae. d) Left, overview image of several microglia (*Hexb*^TdTom^, magenta) and myelin sheaths (AAV-MBP:mem-EGFP, green). Right, zoom of the white box showing several still images of time lapse recording acquired at 10 min intervals. Yellow arrowhead indicates the position of the microglia associated tMB connected to a 16 µm log internode. White arrow indicates the longest observed stretch of the myelin tether of 44 µm at 50 min. e) Quantification of tether stretching velocities (N = 2 tMB, n = 14 push, 16 pull events). f) Vectorial analysis of tMB movement in relation to microglial process position (N = 2 tMB, n = 30 events). g) Patch clamp recordings in voltage-clamp showing increased resistance from 12.9 to ∼70 MΩ during myelin sealing, stretching and tether rupture. h) 2P images of a tMB pulled with the patch pipette. Moving the pipette out of the field of view (>200 µm from the internode) caused the tether to spontaneously rupture. i) 3D rendered image of the pulled tMB.

Next we combined AAV-Mbp:memEGFP labelling with slice cultures prepared from *Hexb*^TdTom^ transgenic reporter mice that express a TdTomato reporter in microglia (*31*). To monitor microglia-myelin interactions, we performed 2-photon imaging in DIV 14-21 slice cultures and recorded a combined volume of 0.05 mm^3^ of tissue for 25.3 hours at 5-20 min intervals. Time-lapse imaging revealed that microglia established persistent contacts with MBs and in some cases, actively stretched them away from the host myelin sheaths forming thin myelin tethers (**Fig. 6d, Suppl. Movie 6**). Microglia processes containing tMBs frequently extended and retracted when pulling the tMBs towards the microglial cell body (**Fig. 6e, Suppl. Movie 6**). We were able to record two of these interactions, during which microglia stretched tMBs by a single microglial process, with tether stretching velocities ranging from 0.05 to 2.09 µm/min (on average 0.59 µm/min, n = 30 events in N=2 tMBs; **Fig. 6e**). The maximum observed tMB stretching was 44 µm from its associated 16-µm-long myelin sheath. Vectorial correlation indicated that directional movement of tMBs followed the trajectory of the associated microglial process suggesting these tMB are indeed actively pulled by microglia processes (**Fig. 6f**). Only very occasionally, tethers broke within the time frame of imaging, resulting in full phagocytosis of the MB by microglia (**Suppl. Movie 6**). The low frequency of tMBs in DIV14–21 cortical slice cultures mirrors their rarity in 6-month-old optic nerves and suggests an age-dependent increase in the prevalence of these myelin–microglia interactions.

### Elasticity of tMBs by Pipette Pulling

To test the basic mechanism of our ‘pulling model’ directly, we probed the ability to slowly mechanically pull off myelin from a mature sheath in the slice by using a patch pipette, controlled by a micromanipulator. Patch pipettes with small open tips (12-14 MΩ resistance) were attached to a single internode (n = 3 internodes pulled, **Fig. 6g**). After forming a high-resistance seal the pipette was slowly retracted (**Fig. 6g,h**) with a speed in line with observed maximal microglial pulling velocities (υ ≍ 2 µm/min). Measured resistance during mechanical myelin pulling demonstrated adherence of myelin membrane to the pipette throughout the pulling process (66 MΩ) (**Fig. 6g**). EGFP imaging revealed thin, highly elastic myelin tethers that could be stretched well over 200 µm before breaking (**Fig. 6h, i, Suppl. Fig. 3**). After tether breakage, the myelin sheaths around axons showed remaining ‘tether tails’, reminiscent of ‘blind tether endings’ protruding from phagocytes or myelin sheaths that we observed in our FIB SEM analysis, and myelin debris could be seen inside the pipette (**Fig. 6h**).

Collectively, live imaging, together with ultrastructural analyses and 3D reconstructions, support a working model in which microglia and astrocytes contribute to physiological myelin turnover by actively pulling spherical myelin fragments from the outer layers of sheaths via a trogocytosis-like mechanism **(Fig. 7a)**. Fusion profiles, loop-like membrane bends, and thinned tethers observed in our EM datasets suggest that tension within the tether brings the innermost lamellae into apposition, promoting their sequential fusion, while separating along the intra-period line (IPL). This sequence progressively weakens the tether until it ruptures. This model can explain how the continuity of the original myelin spiral around the axon is preserved, as apposing lamellae fuse with their correct counterparts during tether thinning **(Fig. 7b)**.

**Fig. 7.**
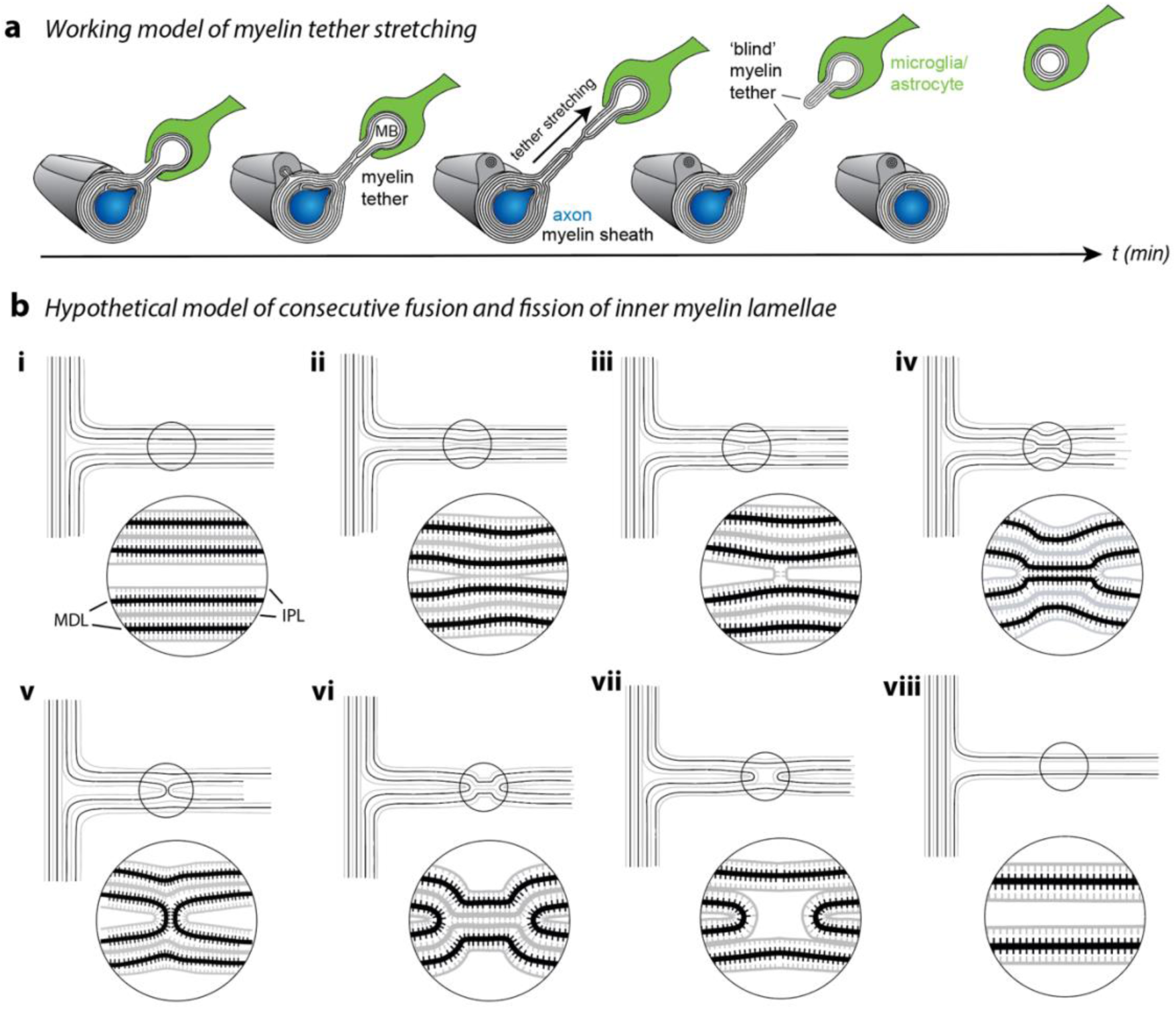
Working model of MB removal from myelin sheaths by myelin tether stretching. Schematic model of trogocytosis by microglia and astrocytes, that turnover myelin. The depicted mechanism proposes that trogocytosis involves (a) stretching of tethered MBs and (b) lateral tension at the outer edges of the pulled myelin tether that forces ordered consecutive delamination at the IPL and intermembrane fusion and fission events of the respective corresponding innermost tether lamellae. This working model can explain how residual myelin sheaths remain structurally organized and intact after removal of a tMBs from an internode, which equally presents with well structured multiple layers of myelin.

## Discussion

We have disovered a novel type of cellular interaction mechanism in the mammalian CNS, by which microglia and astrocytes engage in the turnover of mature myelin, which is made and maintained by oligodendrocytes. The ordered tension-driven excision of multilamellar tMBs directly from myelin sheaths forming a tethered structure that connects the two cell types, is evident in young mouse brains as revealed by time-lapse imaging but becomes more frequent with aging as demonstrated by electron microscopy analyses. The mechanism is related to trogocytosis in other cellular systems, a process in which one cell removes and internalizes small portions of the plasma membrane and membrane-associated proteins from another cell during direct cell–cell contact, with both cells staying alive (*32*).

This aspect of myelin membrane turnover is distinct from other reported mechanisms of debris clearance, such as the engulfment of entire myelin sheaths by microglia in zebrafish larvae (*17*), or the myelin fragmentation into the extracellular space with subsequent clearance by microglia/macrophages (*20–22, 33*), respectively. It also differs from the invasion of myelin sheaths by macrophages/microglia that strip off individual lamellae or entire unravelling sheaths in inflammatory demyelinating disease models (*9–11*). The different behaviour of microglia towards myelin and myelin debris in demyelinating diseases may reflect the duration and degree of myelin damage causing distinct microglial activation states. In agreement with this hypothesis, white-matter associated microglia (WAMs) in the normally aging CNS (*23*) exhibit gene expression profiles that are overlapping but distinct from disease-associated microglia (DAMs) in demyelinating disorders, such as MS, or in Alzheimer’s disease (*34–41*).

Indeed, demyelinating disease models, such as for Multiple Sclerosis or virus-induced demyelination, are characterized by a vast destruction of entire myelin sheaths accompanied by oligodendrocyte death (*42, 43*). Since myelin debris is known to contain inflammatory mediators, neurotoxic lipids and growth inhibiting signaling proteins that interfere with remyelination (*44–46*), efficient myelin clearance in these conditions is critical by sacrificing insulation to contain damage and inflammation as a prerequisite for successful re-myelination. In the context of healthy brain aging and myelin turnover, however, the barriers to microglial invasion may be better preserved, necessitating a more refined mechanism of quantal myelin turnover as suggested by the phagocyte-mediated stretching of tMBs, which preserves the layering and periodicity of the remaining internode.

Our observations are reminiscent of tension-induced trogocytosis in the nervous system that has previously been observed in dendritic spine remodeling, with microglia engulfing spines and physically pulling and stretching spine head filopodia from dendritic shafts by process retraction (*29, 47*). Also actin-bundle containing macrophage filopodia act as cellular tentacles *in vitro*, pulling particles for phagocytic uptake (*48, 49*). Macrophage filipodia can exert retraction forces of several hundred piconewtons over large distances (>10µm) (*49*), with pulling velocities of 40-600 nm/s (*48*). It has been suggested that such a pulling behavior of microglia/macrophages reflects a function of immune cells to mechanically detach pathogens that adhere to a cell surface (*48, 49*). Future studies will explore whether the actomyosin cytoskeleton, a key mechanical actuator in microglia morphodynamics (*50*) and an inducer of tension on mitochondria that contributes to fission (*51*) also provides the necessary tension for the pulling of tMBs.

Our proposed model for tension-driven myelin tether stretching and scission emphasizes the need for the consecutive resealing of always the innermost opposing membranes within the tether (Fig. 7). This sequential mechanism would ensure the continuity of the spiral myelin sheaths after tMB extraction, including the cytoplasm-filled channels essential for axon-glial metabolic support (*52, 53*). Membranes can fuse when brought into close contact with sufficient force. Typically, specialized membrane fusion machineries involving SNARE proteins and calcium generate the tension needed to overcome bilayer repulsion, facilitating membrane fusion (*54, 55*). Is the mere mechanical pulling of tMBs sufficient to bring the innermost myelin membranes in a tether close enough and to trigger their fusion and fission? We suggest that during mechanical pulling the strongest tractive force will affect the outer membranes of the tether, which then compress the inner layers bringing them into fusion distance. We note that *in vitro* increased membrane tension of a tubular double-membrane system indeed first favours and facilitates the fission of the inner opposing membranes (*56*).

The dense and regular membrane spacing of myelin may favour membrane fusion processes within myelin tethers. Much like a strong ‘zipper’, the EM-visible major dense line (MDL) of myelin is compacted by myelin basic protein (MBP), and comprises a merely 2-3 nm wide cytoplasmic space between the cytoplasmic leaflets of the stacked myelin membranes (*57, 58*). To maintain the overall topology and integrity of the myelin sheath and to prevent opening of the cytoplasmic myelin leaflets during the formation myelin tethers, we suggest that pulling delaminates compact myelin along the 4 nm-wide intraperiod line (IPL) (Fig. 7), that is formed by the outer opposing faces of the plasma membranes in myelin (*58*). Thus, our model relies on stronger MBP-mediated adhesion at the MDL compared to the weaker PLP-mediated adhesion at the IPL, which is supported by null-mutant mice phenotypes. Shiverer mice lacking MBP show no MDL compaction, highlighting MBP’s crucial adhesive role (*59*). Conversely, PLP-null mutants and PLP-deficient rats exhibit even tighter IPL compression, suggesting PLP rather acts as a structural strut in extracellular membrane apposition (*60–62*), which mechanical stress may overcome.

Recent studies suggest that also astrocytes participate in phagocytosing myelin debris under pathological conditions (*14, 46*). Our present findings of MBs and tMBs by 3D FIB SEM analyses, as well as PLP-positive lysosomes by immuno-EM, suggest that astrocytes assist in myelin turnover also in the uninjured, aging CNS by similar mechanisms as microglia, i.e. by trogocytosis of tethered MBs. During metamorphosis in *Xenopus laevis* astrocytes are the primary myelin phagocytes in the optic nerve, engulfing myelin whorls still connected to the underlying axon (*63*), reminiscent of tMBs observed in our study. Here, phagocytosis underlies internode shortening during metamorphosis (*63*). We note that shortened internodes have been observed in the aging CNS (*64, 65*), in MS brains and upon spinal cord injury (*66–68*), but have been mostly attributed to remyelination (*64*). Whether repetitive myelin pulling and trogocytosis of tethered MBs affects internode length remains an open question but appears possible when taking into account the enormous pullable membrane reservoir of the myelin sheath that we demonstrated by probing for the elasticity and stretchability of the myelin sheaths with patch pipettes.

While our patch-clamp experiments were not quantitative, the observed remarkable elasticity of myelin tethers in cortical organotypic slices is consistent with low membrane tension in adult oligodendrocytes in culture (*69–72*). Myelin sheath’s elasticity and stretchability likely depends on myelin integrity, lipid composition and myelin gene expression, which are all altered with aging (*73, 74*). Thus, by combining live myelin imaging with quantitative mechanobiological methods, direct probing of myelin sheath elastic properties could become a valuable tool for studying myelin membrane mechanics under pathological and aging conditions.

In conclusion, our study uncovers a novel mechanism of myelin turnover involving tension-driven trogocytosis of tMBs by microglia and astrocytes. Future experiments will have to investigate the impact of mechanical strain on multilamellar myelin and define whether the initiation of tMB pulling requires localized exposure of canonical ‘eat-me’ or ‘find-me’ signals, such as phosphatidylserine (*33*), components of the complement system, or others, akin to microglial processes in synaptic pruning (*75*). This process, distinct from traditional debris clearance, underscores the dynamic interplay of glial cells in preserving myelin integrity. Insights into myelin membrane mechanics and the role of astrocytes in myelin turnover in the aged CNS offer promising avenues for understanding aging and aging-associated diseases with myelin pathology.

## Methods

### Mice

Mice were bred and raised with 2-5 mice per cage at a temperature of 20-24°C and at a humidity of 45-65%. Mice had access to food and water *ad libitum* and experienced a 12-hour light-dark cycle. All experiments were in accordance with the German Animal Welfare Law (Tierschutzgesetz der Bundesrepublik Deutschland, TierSchG). For procedures of sacrificing wildtype C57Bl/6N mice at ages 6, 20 and 24 mo for the subsequent preparation of brains and optic nerves, all regulations given in TierSchG §4 were followed. Sacrificing mice is not an experiment on animals according to TierSchG §7 Abs. 2 Satz 3. All procedures were supervised by the animal welfare officer and the animal welfare committee for the Max Planck Institute for Multidisciplinary Sciences, Göttingen, Germany. The animal facility at the Max Planck Institute for Multidisciplinary Sciences is registered according to §11 Abs. 1 TierSchG as documented by 33.23-42508-066-§11, dated Jan 31^st^, 2024 at the Lower Saxony State Office for Consumer Protection and Food Safety. All animal procedures for the preparation of organotypic slice cultures were done after approval from the Royal Netherlands Academy of Arts and Sciences (KNAW) Animal Ethics Committee (DEC) and Central Authority for Scientific Procedures on Animals (CCD, license AVD80100202216329), and supervised and approved by the Animal Welfare Body (IvD, NIN21.21.01).

### Human Donors and Sample Collection

Postmortem human optic nerve tissue and electron migrographs of n = 14 healthy control donors were used in total, 7 from the Netherlands Brain Bank (NBB, Amsterdam, the Netherlands) and 7 from the Institute for Forensic Medicine at the School of Medicine, University of Saints Cyril and Methodius (Skopje, North Macedonia). Autopsy procedures were approved by the Ethics Committee of the VU University Medical Center (Amsterdam, The Netherlands), and the Institutional Review Boards of the New York State Psychiatric Institute and the Macedonian Academy of Sciences and Arts, respectively. Informed consent was obtained from the donors during life for the use of tissue and clinical data for research purposes. The following subjects were used for electron microscopic and histological analyses:

**Table 1.** Human samples used in electron microscopic analyses. PMD = postmortem delay in hours (h).

| Subject | Sex | Age (years) | Diagnosis | PMD (h) | Brain region | Provider |
| --- | --- | --- | --- | --- | --- | --- |
| 19-026 | M | 55 | Non-demented control, Euthanasia | 05:05 | Optic nerve | NBB |
| 19-036 | F | 85 | Non-demented control, Metastatic lung cancer | 06:50 | Optic nerve | NBB |
| 19-077 | F | 91 | Non-demented control, Old age | 09:30 | Optic nerve | NBB |
| 2020-052 | F | 102 | Non-demented control, Old age | 04:55 | Optic nerve | NBB |
| 2020-054 | F | 81 | Non-demented control, Lymphoplasmatic Lymphoma (Kahler) | 07:40 | Optic nerve | NBB |
| 2020-58 | F | 98 | Non-demented control, Heart failure | 08:45 | Optic nerve | NBB |
| 2020-071 | F | 79 | Non-demented control | NA | Optic nerve | NBB |
| 12355 - INVNW853PME | M | 61 | Non-demented control | 12:00 | Optic nerve | (76) |
| 12325 - INVAK040YFV | M | 39 | Non-demented control | 15:00 | Optic nerve | (76) |
| 12303 - INVNX302BLE | M | 43 | Non-demented control | 15:00 | Optic nerve | (76) |
| 12297 - INVHY646KVZ | F | 57 | Non-demented control | 15:00 | Optic nerve | (76) |
| 12328 - INVWT601YLK | F | 57 | Non-demented control | 35:00 | Optic nerve | (76) |
| 12357 - INVPD269ZJQ | M | 58 | Non-demented control | 18:00 | Optic nerve | (76) |
| 12347 - INVVH454BEE | F | 60 | Non-demented control | NA | Optic nerve | (76) |

### Electron Microscopy

#### Chemical Fixation

Human optic nerve samples from the Institute for Forensic Medicine at the School of Medicine, University of Saints Cyril and Methodius (Skopje, North Macedonia) were processed as described previously (*76*). Dissected optic nerves of mice at ages 6, 20 and 24 mo and human optic nerves obtained from the NBB (The Netherlands, Amsterdam) were immersion fixed in phosphate buffer with 2.5% glutaraldehyde, 4% formaldehyde and 0.5 % NaCl and processed for transmission electron microscopy as described (*77*) or prepared with a modified rOTO-protocol, as described previously (*78*). In brief, the samples were rinsed three times for 15 min with 0.1 M phosphate buffer at 4 °C. Samples were post-fixed for 3 h with 2% OsO_4_ (Science Services, Munich, Germany) and 1.5% K_3_Fe(CN)_6_ at 4 °C and rinsed three times with H_2_O for 15 min each at 4 °C. They were further contrasted with 1% thiocarbohydrazide 1 h at room temperature, followed by three times 15 min rinsing with water and 1.5 h incubation with 2% OsO_4_. After several washes with H_2_O the samples were *en bloc* stained with 2% uranyl acetate overnight at 4 °C. The next day the nerve samples were rinsed three times for 15 min with H_2_O and dehydrated through a series of ascending concentrations of acetone for 15 min each (30%, 50%, 75%, 90%, 3× 100%). The samples were incubated with increasing concentrations of resin (2:1, 1:1, 1:2) for 2 h each and left in 90% Durcupan over night without component D. The next day the samples were incubated with 100% Durcupan (all components) for 4.5 h and polymerized for 48 h at 60 °C. For conventional fixation of cortical organotypic slice cultures, cultures were fixed for 1 hr with freshly made 4% paraformaldehyde (Sigma-Aldrich) + 2.5% glutaraldehyde (aqueous solution, Electron Microscopy Sciences (EMS)) in OSC medium. Slices were then transferred to fresh fixative of 4% paraformaldehyde + 2.5% glutaraldehyde in 0.1 M cacodylate buffer (aqueous solution, EMS) and stored at 4° C for 48 hrs. After fixation, samples were washed in 0.1M cacodylate buffer (aqueous solution, EMS) and postfixed with 1% osmium tetroxide (aqueous solution, EMS), 1.5% potassium ferrocyanide (EMS) in 0.1M cacodylate buffer (EMS) for 45 min. Samples were then washed with MilliQ and dehydrated with increasing concentrations of EtOH; twice with 50% and once in 70%, 80%, 90% for 15 min each, and twice in 100% EtOH for 20 min each. Samples were then infiltrated with 1:1 EtOH-Epon mixture for 1 hr followed by 1 hr in a 1:3 EtOH-Epon mixture. Finaly, samples were placed in fresh Epon (Embed 812, EMS) and stored for 24 hrs at 37 ° C, and polymerized at 65°C for 72 hrs.

#### High-Pressure Freezing and Freeze Substitution

High-pressure freezing and freeze substitution was carried out as described (*77*). In brief, mice were sacrificed by cervical dislocation at ages 6 and 24 mo and optic nerves were dissected, cut and high-pressure frozen using 20% Polyvinylpyrrolidone (PVP, Sigma) in PBS as a filler and the HPM100 high-pressure freezer (Leica, Vienna, Austria). Freeze substitution was carried out in a Leica AFSII (Leica, Vienna, Austria) following this protocol: 0.1% tannic acid in acetone was incubated at −90° C for 100 h followed by three acetone rinses for 30 min each. 2% OsO4, 0.1% uranyl acetate in acetone was used for contrasting the samples (7 h at −90°C). The temperature was automatically raised to −20°C within 14 h, kept there for 16 h and raised to 4°C within 2.5 h. OsO4 was removed by washing four times with acetone for 30 min. The nerves were then rinsed with acetone for 1 h at room temperature. The samples were incubated with increasing concentrations of Epon (Serva, Heidelberg, Germany) (2:1, 1:1, 1:2) for 2 h each and left in 90% Epon overnight. The next day the samples were incubated with 100% Epon for 4.5 h and polymerized for 48 h at 60°C.

#### TEM imaging

Ultrathin sections (80 nm) of embedded human nerve samples were processed at the Institute for Forensic Medicine at the School of Medicine, University of Saints Cyril and Methodius (Skopje, North Macedonia) as described (*76*). Images were acquired at 5000x magnification at random ROIs with a JEM-1400PLUS electron microscope equipped with a 2048 x 2048 digital camera at a resolution of 0.011 micron/pixel (*76*) and digitally edited with ImageJ version 2.1.0. Ultrathin sections of embedded optic nerve samples of mice and human optic nerve samples obtained from the NBB (Amsterdam, The Netherlands) were cut using an ultramicrotome (RMC PowerTome PT-PC, Science Services, Munich, Germany) with a diamond knife (Ultra 35°, 3.0 mm, Diatome, Biel, Switzerland). Ultra-thin sections (50 nm) were collected and transferred onto 100 mesh hexagonal copper grids (Science Services, Munich, Germany), which were coated with formvar. Samples were imaged with a LEO EM912 Omega electron microscope (Carl Zeiss Microscopy GmbH, Oberkochen, Germany) and an on-axis 2048 x 2048 2 k CCD camera (TRS, Moorenweis, Germany) using iTEM (Olympus, Tokyo, Japan) and images were digitally edited with ImageJ version 2.1.0. Images were acquired at 5,000x and 30,000x magnification at random ROIs. In digital images of high pressure frozen optic nerve samples, axon calibers were calculated by measuring the area of the axon and using the formula 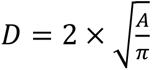 and the number of myelin lamellae was quantified manually. Myelin outfoldings were defined in 2D as finger- or comma-shaped protrusions of compact myelin that included the inner tongue and extended outward. Outfoldings were included into the analysis, when at least as long as the diameter of the originating axon. Relative outfolding area in mouse optic nerves was quantified by placing a grid of 5000 intersections on the field of view (FOV), counting the number of intersections that fall on an outfolding and dividing it by 5000. In humans, outfolding area was manually marked and measured, and then divided by the total FOV area. MBs in 2D analyses were defined as multilamellar, oval myelin structures with visible periodicity and not obviously connected to any surrounding myelin sheaths. A total of 10 FOVs with an area of 480 µm^2^ was quantified per mouse. A total of 24 FOVs with an area of 412 µm^2^ was quantified per human. In part, TEM micrograph datasets of human samples were previously used (*76*), as the basis to reconstruct and analyze features independent from those assessed in the present work. Embedded samples of cortical organotypic slice cultures were cut using a 45° diamond knife (Diatome, Biel, Switzerland) on a Leica Ultracut UCT (Leica Microsystems GmbH, Wetzlar, Germany) to obtain 60 nm thick sections. Sections were placed on formvar covered copper grids (EMS) and post-stained with uranyl acetate (3.5% in ddH_2_O) for 20 min and lead citrate (EMS) for 2 min. A Talos L120C microscope at 120kV with a Ceta 16M camera was used to acquire images.

#### Immunoelectron microscopy

Immunogold labeling on cryosections of cross-sectioned optic nerves from C57Bl/6N mice at ages 6 and 24 mo and electron microscopy was performed as previously described (*77*). In brief, freshly dissected murine optic nerves were fixed in 4% formaldehyde and 0.2 % glutaraldehyde in 0.1 M phosphate buffer containing 0.5% NaCl. Samples were stored in 1% formaldehyde at 4°C. To prepare the nerves for ultrathin cryosectioning, pieces of the nerves were embedded in tiny blocks of 10% gelatin, infiltrated overnight with 2.3 M sucrose in 0.1 M phosphate buffer and then mounted on aluminium pins for ultramicrotomy and frozen in liquid nitrogen. Ultrathin cryosections were prepared with a UC6 cryo-ultramicrotome (Leica Microsystems, Wetzlar, Germany) using a 35° cryo-immuno-diamond knife (Diatome, Biel, Switzerland). For immunolabelling, sections were incubated with an antibody directed against PLP (polyclonal rabbit, home-made (A431, (*79*), 1:250) and subsequently with protein A-gold (10nm) obtained from the Cell Microscopy Center, Department of Cell Biology, University Medical Center Utrecht, The Netherlands. Next, for double immunolabeling, the sections were incubated with antibodies against GFAP (monoclonal mouse, Novocastra NVL-GFAP-GA5, RRID: AB_563739, 1:250) and protein A-gold (15 nm). The sections were then analyzed using a LEO EM912AB (Carl Zeiss Microscopy GmbH, Oberkochen, Germany), and digital micrographs were taken using a 2048x2048 CCD camera (TRS, Moorenweis, Germany).

#### Sample Mounting, FIB-SEM, data acquisition and image analysis

Samples were trimmed with a 90° diamond trimming knife (Diatome AG, Biel, Switzerland) and positioned on a SEM-stub (Science Services GmbH, Pin 12.7 mm x 3.1 mm) using silver conductive resin (EPO-TEK EE 129-4) and polymerized at 60° overnight. Samples were coated with a 10 nm platinum layer using the sputter coating machine EM ACE600 (Leica) at 35 mA current. Samples were placed into the Crossbeam 540 focused ion beam scanning electron microscope (Carl Zeiss Microscopy GmbH, Oberkochen, Germany). To ensure even milling and to protect the surface, a 300 nm platinum layer was deposited on top of the region of interest. Samples were exposed with a 15 nA current and a 7 nA current was used to polish the surface. Datasets were acquired at 1.5 kV using the energy-selective backscatter detector (ESB) in analytical mode with a grid voltage of 450 V and a current of 1000 pA. Using a pixel size of 2 nm × 2 nm and a slicing thickness of 50 nm, a volume of ≥15 μm × ≥ 15 μm × ≥ 197 μm was acquired for the 20 mo old sample and a volume of ≥26 μm × ≥ 26μm × ≥ 75 µm for the 6 mo old sample. All following post-processing steps were performed using imageJ/Fiji (*80*): The images were aligned using the SIFT algorithm, cropped and inverted. They were smoothed using a Gaussian blur (sigma 1) and a local contrast enhancement was applied (CLAHE: blocksize 127, histogram bins 256, maximum slope 1.5). Axons, myelin sheaths, myelin outfoldings, tMBs, astrocytes and microglia were manually segmented in IMOD on datasets that were binned by 5 in x/y for the dataset 20mo and 3 in x/y for the dataset 6mo. The number of MBs was quantified in 1200 individual, consecutive 2D images of the 3D FIB-SEM stack at age 20 mo, blinded to their 3D appearance. Afterwards the 3D identity of each MB^2D^ was checked: (a) free and unconnected to any other myelin structure, (b) connected by a tether of thinner diameter than the MB or (c) any other form of myelin protrusions or deformity. Several individual MBs^2D^ can therefore belong to the same 3D structure. MBs in the 2D images were defined as multilamellar, spherical or oval myelin structures with often normal lamella periodicity and not connected to a myelin sheath.

#### SEM imaging

Human optic nerves, embedded with the rOTO protocol, were sectioned using an UC7 ultramicrotome (Leica Microsystems GmbH, Wetzlar, Germany) with an array tomography ATS diamond knife (Diatome, Biel, Switzerland). Sections of 0.2 µm thickness were collected on carbon-coated 10 mm x 10 mm silicon wafer (Science Services, Munich, Germany). After coating the sections again with 5 nm carbon with a Leica ACE600 Sputter Coater (Leica Microsystems GmbH, Wetzlar, Germany), SEM image tiles were obtained with a Crossbeam 540 focused ion beam scanning electron microscope (Carl Zeiss Microscopy GmbH, Oberkochen, Germany) at 15 nm pixel size at 3.0 kV with a probe current of 1.0 nA and a dwell time of 10 µs with the BSD1 backscatter detector at a working distance of 6 mm using the ATLAS5 software (Mosaic tile scan, Fibics/Carl Zeiss Microscopy GmbH, Oberkochen, Germany).

### Luxol Fast Blue

To stain myelin and myelinated axons, paraffin-embeded human optic nerve samples were used that were previously generated (*81*) as the basis to analyze features independent from those assessed in the present work. 8 µm paraffin sections were deparaffinzed and rehydrated in a xylene and ethanol series, then stained in 0.1% Luxol Fast Blue (Sigma-Aldrich), 95% ethyl alcohol, and 0.5% glacial acetic acid (Merck) at 60°C o.N. After washing in 95% alcohol and distilled water, they were differentiated in 0.05% lithium carbonate in distilled water for 30 sec, rinsed in distilled water for 1 min, and cleared in 70% alcohol for until only white matter remained visible. Sections were counterstained with Mayer’s hematoxylin (Sigma-Aldrich) for 1 min, dehydrated and mounted with Permount (Thermo-Fisher Scientific).

### Immunohistochemistry

Mice at the ages of 6 and 24 mo were injected intraperitoneally with Avertin and lethal overdose was ensured prior to transcardial perfusion with Hank’s buffered salt solution (HBSS, Gibco, Thermo Fisher Scientific) followed by 4% Paraformaldehyde (PFA) in 0.1 M phosphate buffer. Brains were dissected, fixed overnight in 4% PFA in 0.1 M phosphate buffer, paraffinized using a Microm STP 120 Spin Tissue Processor (Thermo Scientific) and then embedded in paraffin blocks on a HistoStar embedding workstation (Epredia). Brains were cut into 5 µm sections on a RM2155 (Leica) rotary microtome and mounted on adhesive microscope slides (Histobond). Slices were deparaffinized by incubation in xylene followed by a descending ethanol series. Heat-induced epitope-retrieval was performed in sodium citrate buffer (0.01 M, pH 6.0, 10 min in a 400-Watt microwave with a 20 min cooldown period). Slides were blocked for 1h at RT in 20% goat serum and 2% BSA in PBS and primary antibodies were diluted in 2% BSA in PBS. We used rabbit anti-MBP, 1:100 (*25*); mouse anti-GFAP, 1:200 (Chemicon) and rat anti-Iba1, 1:2000 (Abcam). Primary antibody incubation was done at 4°C overnight and secondary antibody incubation (goat anti-rabbit IgG (H+L) antibody, DyLight 800 Conjugated, 1:5000 (Rockland); donkey anti-rat IgG (H+L) A555 (Abcam); donkey anti-mouse IgG (H+L) A555 (Abcam)) at RT together with DAPI (1:10,000) to label cell nuclei for 1h. Finally, slides were washed 3 x 5min in 1x PBS before mounting in embedding medium ProLong Glass (Invitrogen, Thermo Fisher Scientific) and storing at 4°C until imaging. Widefield images of stratum lacunosum moleculare (SLM) were taken using an Axio Imager.Z1 (Zeiss) equipped with a Colibri 7 solid state light source using an ORCA-Fusion CMOS Camera (C14440-20UP, Hamamatsu) using a 40x lense (NA = 1.4), an x/y resolution of 2.4 px/µm2, and a total field size of 1 mm². The number of astrocytes and microglia containing PLP inclusions in the field of view were counted and divided by the total number of astrocytes and microglia, respectively. Cofocal images of the SLM in the CA1 hippocampal region were acquired using the 100x PI APO lense (NA = 1.4) of the Leica TCS SP5 with an optical thickness of 0.713 µm, an x/y resolution of 6.6 µm/px, and a total field size of 436 µm x 572 µm. Myelin inclusion volumes were calculated by respectively measuring connected PLP + GFAP and PLP + Iba1 colocalizing areas across z-slices and multiplying by the optical thickness. Particles <0.2 µm³ were excluded to reduce false positives. For both widefield and confocal microscopy, GFAP and Iba1 immunofluorescence was acquired each at 594 nm and PLP at 633 nm wavelength.

### Cortical Organotypic Slice Cultures

Cortical organotypic slice cultures were prepared from 4-5 day old mouse pups. Briefly, mice were anaesthetized via hypothermia, then sacrificed by decapitation with scissors. The brain was extracted and placed in ice-cold dissection solution consisting of 98% GBSS (in mM) 1.5 CaCl_2_, 0.2 KH_2_PO_4_, 0.3 MgSO_4_, 2.7 NaHCO_3_, 5.0 KCl, 1.0 MgCl_2_, 137 NaCl, 0.85 Na_2_HPO_4_, 5.6 D-glucose), 1% (0.1 M stock) kynurenic acid, and 1% (2.5 M stock glucose), 1% ( 0.1 M stock kynurenic acid), sterile filtered with 0.2 µm filtration flasks, and adjusted to pH 7.2 and 320 mOsm. Under a dissection microscope, brains were cut down the midline with a scalpel and then sectioned via McIlwain Tissue Chopper to obtain 300 µm thick coronal slices. Slices were quickly transferred to hydrophilic PTFE membrane inserts (Merck-Millipore, PICMORG50) in 6-well plates and cultured at 35°C and 5% CO_2_ in culturing medium consisting of 47.75% MEM (Thermo Fisher Scientific # 11575032), 25% HBSS (Thermo Fisher Scientific # 24020133), 50% heat-inactivated horse serum (Thermo Fisher Scientific # 26050088), 2% (2.5 M stock) D-glucose, and 1.25% (1 M stock) HEPES (Sigma-Aldrich H3375), sterile filtered, and adjusted to pH 7.2 and 320 mOsm. Medium was changed 3 x a week with fresh equilibrated medium. AAV-MBP:mem-EGFP was applied directly to slices cultures at DIV 7 to selectively label oligodendrocytes. For immunohistochemistry, cultures were fixed with 4% PFA in PBS for 20 min followed by 3 x 10 min washes in PBS. Slices were then blocked for 2 hrs in PBS containing 0.5% Triton X-100 and 10% normal goat serum. Next, slices were incubated 0.25% Triton X-100 and 5% normal goat serum in PBS containing primary antibodies (Supplementary Table 1), overnight at room temperature while shaking. Slices were again washed 3 x 10 min in PBS. Slices were incubated in PBS containing secondary antibodies for 2 hrs at room temperature while shaking and protected from light. Slices were finally washed in PBS 3 x 10 min before mounting with Vectashield mounting medium containing DAPI (Vector Laboratories #H-200). Stained slices were imaged on a Leica SP8 confocal microscope using a 40x 1.3 NA oil-immersion lens and running LASX (3.5.7). Tile-scans were acquired using sequential scans of individual channels using step sizes of 0.346 µm along the Z-axis and at a 2048x2048 pixel resolution. Images were imported in FIJI (FIJI 64 bit; ImageJ version 1.54p; RRID: SCR_002285) and maximum projection function was used to generate Figure 6d.

### Live Two-Photon Imaging

14-21 DIV cortical organotypic slice cultures were placed in the recording chamber of a two-photon microscope (Femto-3D-RD, Femtonics), perfused with carbogen-bubbled recording solution (in mM: 125 NaCl, 25 NaHCO_3_, 1.25 NaH_2_PO_4_, 3 KCl, 25 D-Glucose, 2 CaCl_2_, 1 MgCl_2_). *Hexb*^TdTom^ microglia and mem-EGFP labelled oligodendrocytes were visualized via a Ti:Sapphire laser (Chameleon Ultra II; Coherent, Inc.) tuned to 980 nm, cells were imaged using a 1.0 NA 20x lens (Olympus) with a voxel size of 0.3 x 0.3 x 1 µm (X/Y/Z) at 5-20 min intervals. Image acquisition took place in MES software (Femtonics Inc., version 6.3.7902). Acquired images were corrected for drift using Fast4DReg plugin for FIJI. The velocity of tMBs pulled by microglial processes were analysed using the TrackMate plugin (*82*). The FIJI line tool was used to measure both the trajectory of tMB and microglial processes between frames and the difference between these two angles was used to correlate tMB displacement with microglial processes movement. Patch pipettes with small pipette tip size (12-14 MΩ) were used to mechanically seal to myelin sheaths with brief suction before retracting the tip of the pipette away from the targeted internode. A representative 3D image of a pipette pulled myelin tether was generated using IMARIS (10.01) (Figure 6i).

### Statistics

All statistics and quantifications were performed blinded. Myelin inclusions in astrocytes and microglia of confocal images were analyzed the following: A linear mixed model (LMM) was fitted on log transformed volume data and controlled for random effects from replicates of the same animal (log(Volume) ∼ Age + (1 | Animal) via restricted maximum likelihood (REML) method using the lme4 package (1.1.37) in R (4.4.1). Estimated marginal means and pairwise comparisons were calculated using the emmeans package (1.8.7). For two sample comparisons Student’s t-test was used and for multiple comparison a multiple t-test with Holm-Šidák correction or two-way ANOVA with Šidák correction, as indicated in the figure legends. For the exception of the LMM, statistics was performed on GraphPad Prism (10.4.0). Levels of significance are indicated as n.s = not significant, *p<0.05, **p<0.01, ***p<0.001

## Data Availability Statement

The data that support the findings of this study are available from the corresponding author upon request. Focussed Ion beam scanning electron microscopy will be deposited to the Electron Microscopy Public Image Archive (EMPIAR).

## Acknowledgements

We thank Ulli Bode, Christos Nardis, Johannes Werner Pauly, and Annnette Fahrenholz for technical assistance, and all personnel of the animal facility at the Max Planck Institute for Multidisciplinary Sciences (MPI-NAT), City Campus for animal husbandry. The authors are also thankful for the support of the Electron Microscopy Centre Amsterdam (EMCA) and like to thank Prof. Dr. med. Martin Kerschensteiner (University of München) for sharing the AAV-Mbp:mem-EGFP plasmid, Dr. Koen Kole for producing the virus and Prof. Dr. Marco Prinz (University of Freiburg) for mice expressing TdTomato via the Hexb promoter (*Hexb*^TdTom^). We also thank Prof. Dr. Inge Huitinga and the Netherlands Brain bank (NBB) for providing post-mortem human optic nerve tissues. This work was financially supported by grants of the German Research Foundation/DFG to SG (SPP1757), WM (Cluster of Excellence and DGF Research Center Nanoscale Microscopy and Molecular Physiology of the Brain and MO 1084/2-1 (FOR 2848)), and KAN (TRR274), who also received support by the Dr. Myriam and Sheldon Adelson Medical Foundation (AMRF) and the European Research Council (Advanced Grant MyeliNANO). M.K. and K.H. acknowledge research support by funding from the Institute of Chemical Immunology (project ICI0000030) and from the Netherlands Institute for Neuroscience (NIN) Friends Foundation.

## Author contributions

ES Conceptualization, Investigation, Data analyses, Writing-original draft, Writing-review & editing

KPH Methodology, Investigation, Data analyses, Writing-review & editing

YK Investigation, Data analyses, Writing-review & editing

CM Investigation, Data analyses, Writing-review & editing

LK Methodology, Writing-review & editing

MT Methodology, Writing-review & editing

UG Investigation, Writing-review & editing

SM Data analyses, Writing-review & editing

SL Investigation, Writing-review & editing

BS Methodology, Writing-review & editing

TR Methodology, Writing-review & editing

PAJ Unpublished materials, Writing-review & editing

AJD Unpublished materials, Writing-review & editing

IH Unpublished materials, Writing-review & editing

MHPK Supervision, Writing-review & editing

AS Methodology, Investigation, Writing-review & editing

WM Methodology, Supervision, Funding, Writing-review & editing

KAN Funding, Writing-review & editing

SG Conceptualization, Funding, Supervision, Investigation, Data analyses, Writing-original draft, Writing-review & editing

