## Supplementary Files for "Tether-mediated extraction of myelinoid bodies by microglia and astrocytes can maintain myelin integrity"

### Supplementary Material

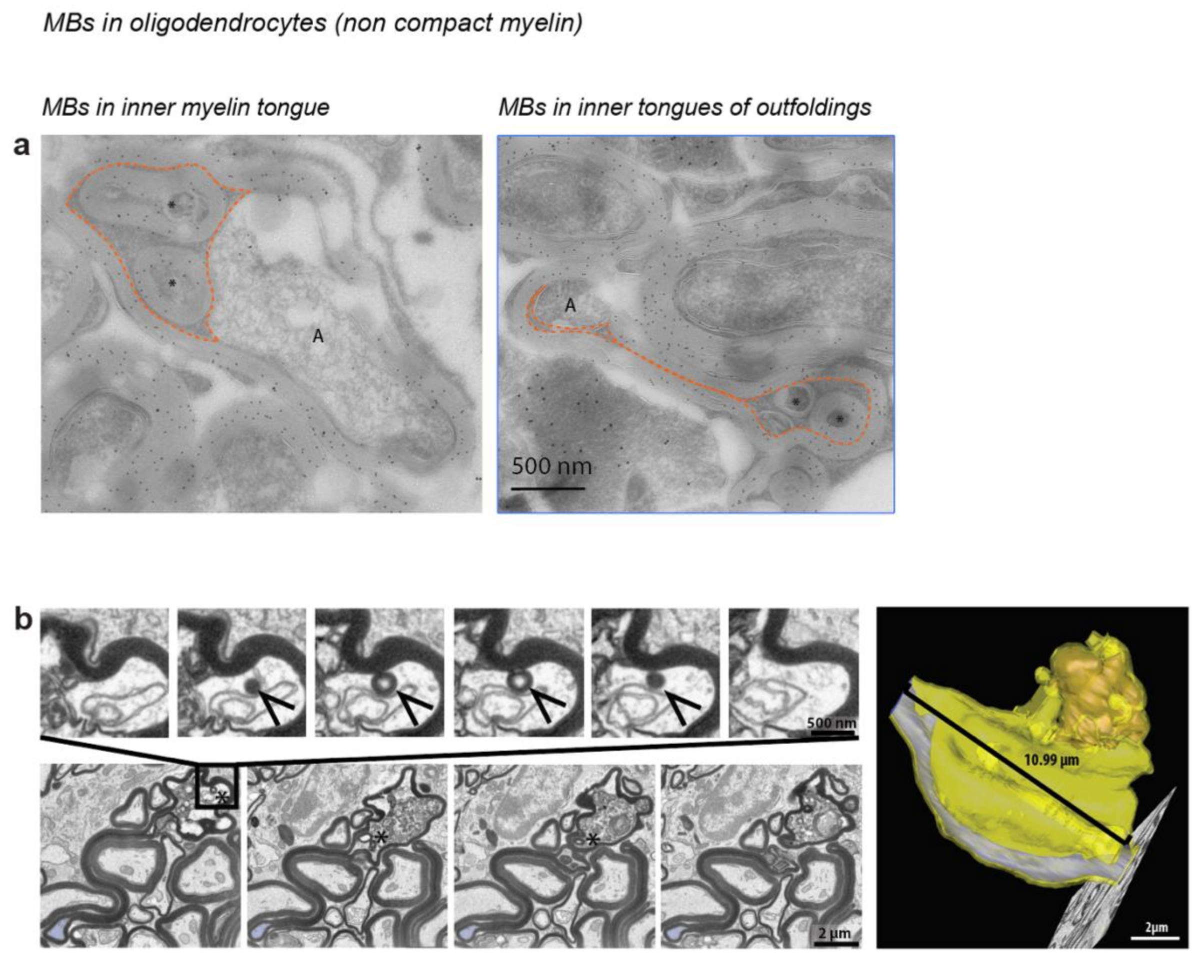


**Supplementary Fig. 1 MBs in non-compact myelin**

**a)** Colloidal immunogold-EM labelling on ultrathin optic nerve sections of wildtype mice at age 24 mo. MBs (co-labelled for PLP by 10 nm gold-particles) localize in non-compact inner tongues of myelin sheaths (left, orange dashed line) and in swollen inner tongues of myelin outfoldings (right, orange dashed line). GFAP is visualized by 15 nm gold-particles. **b)** 3D reconstruction (right) of the tip of an outfolding, which is filled with vesicular debris and few MBs (bottom left, single FIB-SEM images from reconstruction; top left, higher magnification of an MB (arrowhead) that constricts into the swollen inner tongue of the tip of the myelin outfolding), asterisks, MBs; A, axon.

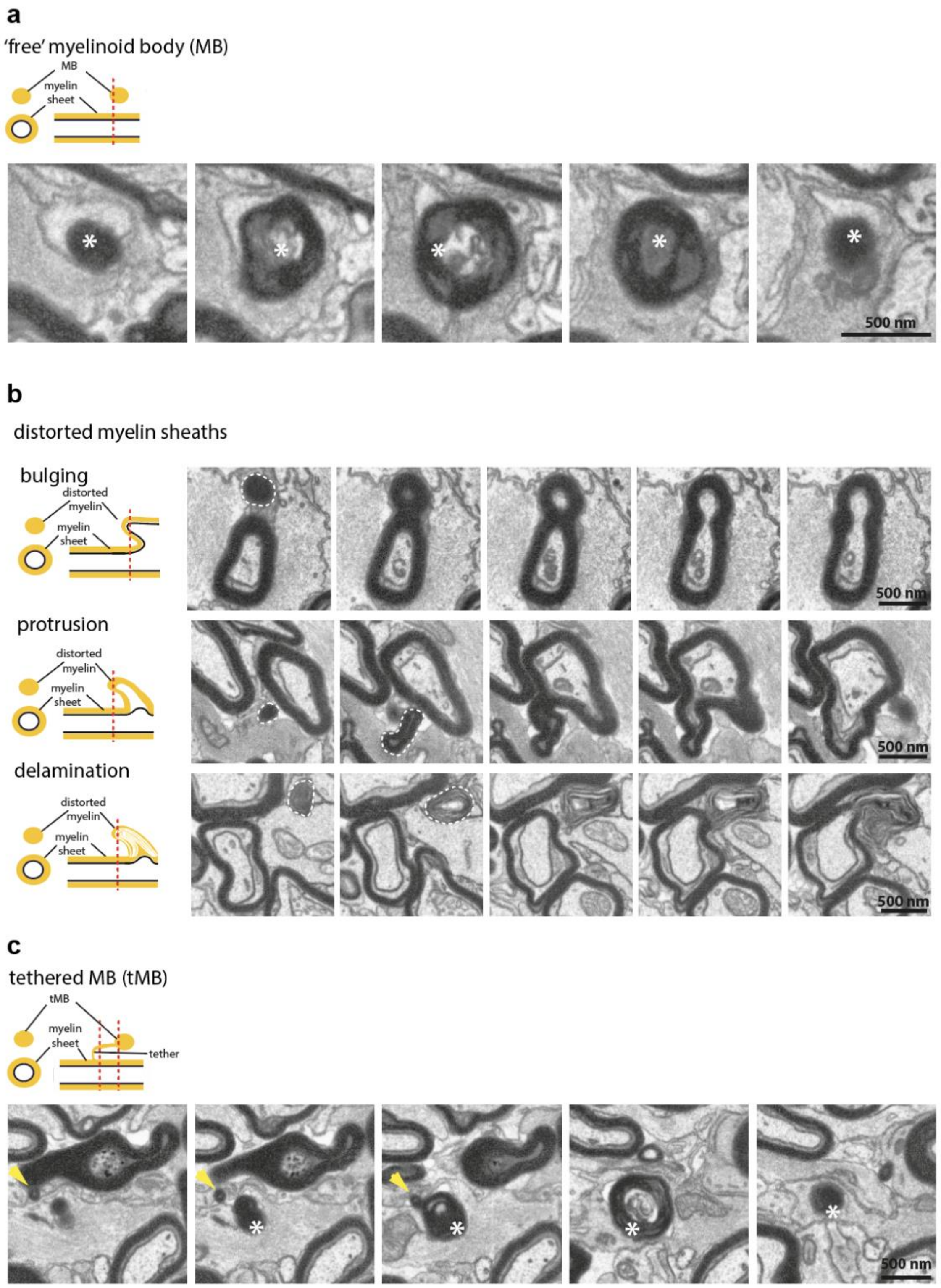


**Supplementary Fig. 2 Classification of 2D myelin bodies**

Several different structures can appear as MBs in 2D. Based on their 3D identity, $MBs^{2D}$ can be separated into three distinct classes: **a)** 'Free' MBs (asterisks) that are not connected to the myelin sheet in 3D in any way, **b)** axonal outbulgings, protrusions, as well as delaminated myelin, that appear as disconnected MBs in 2D, or **c)** MBs (asterisks) that are still connected to the myelin sheet via a thin tether (arrowhead) in 3D (tMBs). Here both the spheroid of the tMB and the tether itself can appear as MB in 2D.

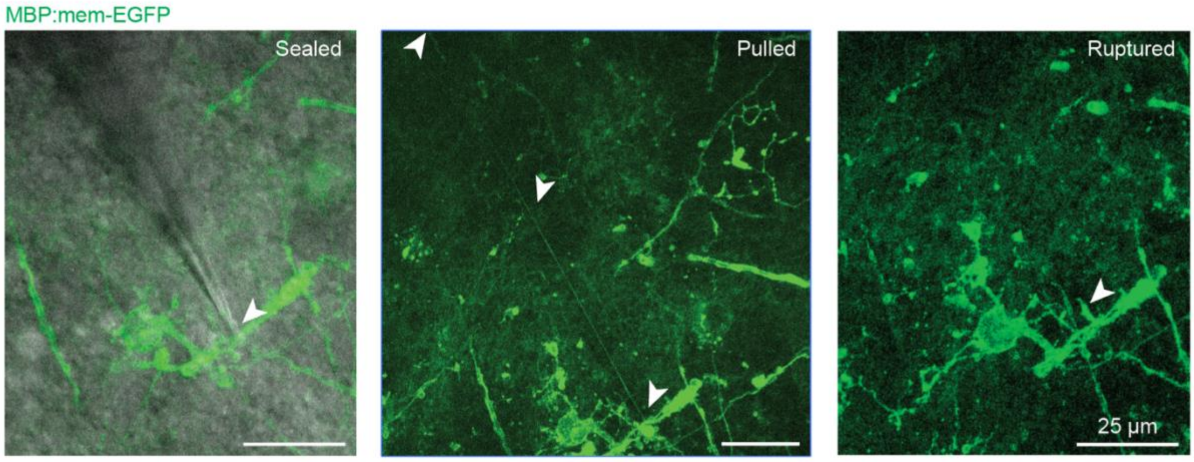


**Supplementary Fig. 3 Elasticity of tMBs by pipette pulling**

2P images of a tMB pulled with a patch pipette. Left, pipette is attached to the myelin sheath. Middle, tether stretched to 228 µm via patch pipette. White arrowhead indicate the tether (including the beginning and end). Right, image acquired after pipette was retracted out of the field of view which caused spontaneous rupture of the tether. The tether tail is indicated by a white arrowhead.

#### Supplementary Movie 1

**3D reconstruction of myelin outfoldings in an aged mouse optic nerve.** FIB-SEM analysis of a 20 mo old wild-type mouse optic nerve shows that myelin outfoldings represent sheets in 3D. Three axon segments are pseudo-colored in blue and their myelin sheaths and outfoldings in yellow for compact myelin and orange for non-compact myelin. Two MBs are connected to either a myelin sheath or an outfolding, respectively by short myelin tethers. Related to Fig. 3a.

#### Supplementary Movie 2

3D reconstruction of aberrant myelin structures in an aged mouse optic nerve. FIB-SEM of a volume of 237µm$^3$ of a 20 mo old wild-type mouse optic nerve shows that aberrant myelin ultrastructures are abundant during aging. MBs are preferentially still connected to myelin sheaths by thin, long myelin tethers (=tMBs) and are often engulfed by astrocytes or microglia. Blind ending tethers indicate tether rupture. Axons are pseudocolored in blue and their myelin in yellow. Astrocytes are pseudocolored in green and microglia in cyan. Related to Fig. 3c.

**Supplementary Movie 3**

3D reconstruction of a tMB engulfed by microglia in an aged (20 mo) mouse optic nerve. The spheroid of the tMB is fully surrounded by the microglia with the exception of the tether that is still connected to the myelin sheath. Degraded myelin debris inside the microglia indicates it is actively involved in myelin degradation. Movie shows EM images with super-imposed pseudo-colored 3D reconstructions. Axons are pseudocolored in blue, myelin in yellow, microglia in cyan, myelin debris in white and astrocyte in green. Related to Fig. 3d.

**Supplementary Movie 4**

3D reconstruction of an astrocyte process (green) in an aged mouse optic nerve (20m) that has engulfed the spheroid of a tMB. Movie shows FIB-SEM images with super-imposed pseudo-colored 3D reconstructions (axon segment, blue; myelin outfolding and tMB, yellow;). Related to Fig. 4c.

**Supplementary Movie 5**

3D reconstruction (mouse optic nerve; age, 6 mo) of a tMB (yellow) engulfed by an astroglial process (green) that is connected via a short myelin tether to a myelin sheath **(**yellow).

**Supplementary Movie 6**

Two-photon imaging of microglia (*Hexb*$^{Tdtom}$) and oligodendrocytes (MBP:mem-EGFP) in cortical organotypic slice cultures. Example #1: Prolonged microglia interaction with tMB. Example #2: Microglia successfully breaks a tMB, which is rapidly leaving the microscopic field. Yellow arrow denotes the moment the tether is broken. Example #3: Prolonged microglia interaction with tMB on the edge of field of view. Example #4: Microglia-myelin interaction.

### Supplementary Table 1

| Antibody | Host | Dilution | Manufacturer | Cat. number | RRID |
|---|---|---|---|---|---|
| Anti-GFP | Chicken | 1:500 | Abcam | ab13970 | AB_300798 |
| Anti-IBA1 | Guinea pig | 1:500 | SySy | 234 308 | AB_2924932 |
| Anti-MBP | Rat | 1:200 | Millipore | MAB386 | AB_94975 |
| Anti-Chicken-Alexa-488 | Goat | 1:500 | Thermo Fisher | A11039 | AB_142924 |
| Anti-Guinea pig-Alexa-594 | Goat | 1:500 | Thermo Fisher | A11076 | AB_141930 |
| Anti-Rat-Alexa-647 | Goat | 1:500 | Thermo Fisher | A21247 | AB_141778 |
